# The deubiquitinating activity of Middle East respiratory syndrome coronavirus papain-like protease delays the innate immune response and enhances virulence in a mouse model

**DOI:** 10.1101/751578

**Authors:** Robert C. M. Knaap, Raúl Fernández-Delgado, Tim J. Dalebout, Nadia Oreshkova, Peter J. Bredenbeek, Luis Enjuanes, Isabel Sola, Eric J. Snijder, Marjolein Kikkert

## Abstract

Middle East respiratory syndrome coronavirus (MERS-CoV) continues to cause zoonotic infections and serious disease, primarily in the Arabian Peninsula, due to repeated spill-over from dromedary camels and subsequent nosocomial transmission. Approved MERS vaccines for use in animals or humans are not currently available. MERS-CoV replication requires the virus-encoded papain-like protease (PL^pro^) to cleave multiple sites in the viral replicase polyproteins, thereby releasing functional non-structural proteins. Additionally, PL^pro^ is a deubiquitinating enzyme (DUB) that can remove ubiquitin(-like) moieties from substrates, presumably to counteract host antiviral responses. In previous work, we determined the crystal structure of MERS-CoV PL^pro^ in complex with ubiquitin, facilitating the design of PL^pro^ mutations that impair DUB activity without affecting viral polyprotein cleavage. Here, we introduced these DUB-inactivating mutations into the viral genome and examined their impact on MERS-CoV infection both in cell culture and in a lethal mouse model. Although overall replication of DUB-negative and wild-type (wt) recombinant MERS-CoV was comparable in multiple cell lines, infection with DUB-negative virus markedly increased mRNA levels for interferon (IFN)-β and IFN-stimulated genes. Moreover, compared to a wt virus infection, the survival rate was significantly increased when DUB-negative MERS-CoV was used to infect transgenic mice expressing a human MERS-CoV receptor. Interestingly, DUB-negative and wt MERS-CoV replicated to the same titers in lungs of infected mice, but the DUB-negative virus was cleared faster, likely due to the observed accelerated and better-regulated innate immune responses, in contrast to delayed and subsequently excessive responses in wt virus-infected mice. This study provides the first direct evidence that the DUB activity of a coronaviral protease contributes to innate immune evasion and can profoundly enhance virulence in an animal model. Thus, reduction or removal of the innate immune-suppressive DUB activity of PL^pro^s is a promising strategy for coronavirus attenuation in the context of rational vaccine development.

**Author Summary:** Although zoonotic coronaviruses such as Middle East respiratory coronavirus (MERS-CoV) have pandemic potential, therapeutics and vaccines that counteract this public health threat are not currently available. Coronaviruses typically employ multiple strategies to evade the host’s innate immune response, which may enhance clinical disease and/or reduce the efficacy of modified live vaccines. The MERS-CoV-encoded papain-like protease (PL^pro^) is not only crucial for the expression of functional replicase proteins, but has also been postulated to antagonize ubiquitination-dependent steps during the activation of the innate immune response. Here, we report the generation of engineered MERS-CoVs mutants in which PL^pro^’s deubiquitinating (DUB) activity was specifically disrupted without affecting virus viability. In this manner, we could demonstrate that the DUB activity of PL^pro^ suppresses the interferon response in MERS-CoV-infected cells. Strikingly, in the lungs of mice infected with DUB-negative MERS-CoV, innate immune responses were induced at an earlier stage of infection than in wt virus-infected mice. This group also showed a clearly increased survival, indicating that the DUB activity is an important MERS-CoV virulence factor. This proof-of-concept study establishes that the engineering of DUB-negative coronaviruses, which elicit a more effective immune response in the host, is a viable strategy for vaccine development.

## Introduction

The *Coronaviridae* family includes numerous viruses that can cause respiratory and enteric disease in animals or humans. To date, six coronaviruses (CoVs) have been identified that have the ability to infect humans, including two that have caused global public health concerns due to high case-fatality rates. Firstly, in 2003, an outbreak of severe acute respiratory syndrome coronavirus (SARS-CoV) led to about 8000 confirmed human infections with a case-fatality rate of around 10% (1). Fortunately, within 6 months, the epidemic spread of SARS-CoV could be halted by conventional public health-and quarantine measures. Secondly, since 2012, about 2500 laboratory-confirmed cases of human infection with Middle East respiratory syndrome coronavirus (MERS-CoV) have been reported (2). Given the ∼35% MERS case-fatality rate, this implies that the number of MERS-related deaths now exceeds the death toll of the 2003 SARS-CoV outbreak. Although MERS cases were reported from 27 different countries, the vast majority of human infections have occurred in Saudi Arabia where the virus was also first recovered from an infected individual. It was subsequently identified as a previously unknown CoV by genome sequence analysis (3, 4). MERS-CoV appears to be widespread among dromedary camels (5), which shed the virus from their upper respiratory tract (6) and were identified as a reservoir from which repeated spill-over to humans has occurred (7, 8). Nevertheless, bats, which harbour many different CoV species including a variety of SARS- and MERS-like CoVs, are considered to be the most likely wild-life reservoir for both these viruses (9, 10). The MERS problems are further augmented by secondary human infections acquired in household settings or health-care facilities (11). To date, therapeutics nor vaccines to prevent or treat MERS-CoV infections have been approved.

Host organisms have evolved antiviral defense systems, whereas pathogens in turn have adapted to evade these systems or suppress their activation (12). Upon infection, the vertebrate host’s first line of defense is the innate immune response, which is initiated when pathogen-associated molecular patterns, such as double-stranded (ds) RNA, are sensed by pathogen recognition receptors (13). After initial sensing, activation of subsequent signalling cascades leads to nuclear translocation of the transcription factors interferon regulatory factor (IRF) 3, IRF7 and NF-κB, which drives expression of antiviral molecules such as interferons (IFNs) and other pro-inflammatory cytokines. Type I IFNs (IFN-α/β) and type III IFNs (IFN-λ) induce expression of interferon-stimulated genes (ISGs) with the distinction that, due to receptor restrictions, IFN-λ can only induce ISGs in epithelial cells, whereas ISGs induced by type I IFNs can be expressed by virtually any cell type (14, 15). ISGs are antiviral effector proteins that limit virus replication and spread. Moreover, some ISGs can enhance pathogen recognition and thereby innate immune signalling (16). NF-κB-mediated production of cytokines like TNF-α, IL-6, and IL-8, are important for the inflammatory response as well as for attracting and activating immune cells (17). Type I IFNs also play a major role in stimulating expression and production of other immune mediators like IFN-γ and cytokines by diverse innate immune cells. Additionally, antigen presentation by dendritic cells is enhanced and also the survival and activity of B and T cells is modulated by type I IFNs, which underlines the importance of type I IFNs in shaping adaptive immune responses (18, 19). The fine-tuning of innate immune responses is crucial and therefore the activities of signalling molecules in these pathways are tightly regulated by post-translational modifications such as phosphorylation and ubiquitination (20). Ubiquitination is the covalent conjugation of ubiquitin (Ub), an evolutionally conserved protein, via attachment of its C-terminal Gly76 residue to the N-terminal residue or any Lys residue of a protein substrate (21). Conjugation of Ub can also occur to previously conjugated Ub moieties on the N-terminus or at any of the seven Lys residues in Ub itself, leading to the formation of polyUb chains with different topologies and functions (22, 23). Ubiquitination is performed by a cascade of three enzymes: an E1 Ub-activating enzyme, an E2 Ub-conjugating enzyme and an E3 Ub ligase. The reverse reaction, the removal of Ub from substrates, is performed by proteases that are known as deubiquitinating enzymes (DUBs) (24). In the context of the innate immune response, ubiquitination is essential for the activation or induced degradation of many factors in the signalling cascade, while specific cellular DUBs can downregulate the signalling to protect cells from adverse over-reactions of the system (25, 26). In response, many viruses have developed means to evade activation of the immune response by modulating the Ub-system, for example by expressing virus-encoded DUBs (vDUBs) or E3 ligases, or by hijacking or blocking the activity of host DUBs or E3 ligases (27, 28).

Coronaviruses are positive-stranded RNA viruses with large genomes of 26-to 34-kb, which have a highly conserved organization (29). The first two-thirds of their genome encode the replicase polyproteins (pp1a and pp1ab) that yield 15 or 16 non-structural proteins (nsps) following cleavage by internally-encoded proteases. Structural and accessory proteins are expressed from the remaining one-third of the genome, using a nested set of viral subgenomic RNAs that are produced in infected cells (30). Viral RNA synthesis is thought to occur in close association with virus-induced membranous ‘replication organelles’, including double-membrane vesicles that label abundantly for dsRNA (31). Melanoma differentiation-associated gene 5 (MDA5), a cytosolic innate immune sensor, was proven to be crucial for detecting murine CoV replication (32). Despite the production of viral RNA species that can activate the host’s innate immune response, IFNs and pro-inflammatory cytokines are hardly expressed or only appear late in cell culture-based infection models for various CoVs (33-37). In a mouse model for SARS-CoV infection, a delayed and subsequently excessive IFN response was shown to induce severe lung immunopathology, due to the influx of inflammatory monocyte-macrophages leading to elevated levels of cytokines and chemokines (38). Similar pathological observations were made in a mouse model for lethal MERS-CoV infection, which is based on a mouse-adapted virus obtained by serial passaging of the parental virus in mice (39). Coronaviruses employ various strategies to prevent immune detection and actively suppress or delay activation of the immune system, which is thought to contribute to viral virulence and pathogenesis (40). The structural membrane (M) protein and multiple accessory proteins of MERS-CoV have been shown to antagonize type I IFN- as well as NF-κB responses (41-46). Several other mechanisms of interference with the innate immune response were attributed to CoV nsps, specifically nsp1, 3, 15, and 16. Nsp1 of β-CoVs has been shown to alter host antiviral gene expression by selectively targeting mRNAs for degradation (47, 48). The CoV macrodomain in nsp3, which has de-ADP-ribosylation activity, antagonizes innate immune signalling (49, 50). Moreover, inactivation of the nsp15 endoribonuclease led to a severely attenuated virus that strongly activated sensors MDA5, PKR, and 2’,5’-oligoadenylate synthetase (51, 52), suggesting that the enzyme normally serves to prevent activation of the immune response. Inactivation of the MHV nsp16 2’*O*-methyltransferase, which was shown to be pivotal for protection of CoV RNA from recognition by MDA5, enhanced IFN-β expression upon infection (53). In contrast, inactivation of this activity in SARS-CoV did not induce a robust IFN-β response, but it did attenuate the virus in cell culture-based infections and mice (54). Finally, the papain-like protease (PLP) in nsp3 can bind and cleave Ub and ISG15 from substrates, most likely to suppress the antiviral responses (reviewed by (55)). In the case of α-CoVs and lineage A β-CoVs, nsp3 contains two PLPs (PLP1 and PLP2), of which usually one has DUB activity, while the remaining β-CoVs, and also γ- and δ-CoVs, express a single PLP (PL^pro^) with DUB activity. The DUB activity of CoV PLPs was first discovered for SARS-CoV and subsequently PL^pro^ was shown to suppress innate immune signalling pathways upon its ectopic expression (56-58). In similar experimental settings, these features have now been demonstrated for a variety of CoV PLPs, including MERS-CoV PL^pro^ (55, 59, 60).

Disruption of antagonists of the IFN response can be an approach for modified live virus (MLV) vaccine design because such virus mutants will still possess the natural immunogens, while being attenuated-by-design. Due to their reduced capability to evade the innate immune system, they would be anticipated to induce enhanced immune responses. This concept was successfully applied to influenza virus A and B by removing or mutating the NS1 protein, a well-characterized IFN antagonist. This yielded strongly attenuated viruses that provided protective immunity in mice and induced neutralizing antibodies in humans (61-63). More recently, *in vivo* attenuation of MERS-CoV in a mouse model was achieved by deletion of open reading frames (ORFs) 3 to 5, or by inactivation of the nsp16 2’*O*-methyltransferase (45, 64). In our current study, we aimed to eliminate the DUB activity of MERS-CoV PL^pro^ to evaluate its role during infection in cell culture as well as a mouse model, and to determine if a DUB-negative MERS-CoV is attenuated and a potential MLV vaccine candidate. Based on the crystal structure of the MERS-CoV PL^pro^-Ub complex, the Ub-binding site of PL^pro^ was equipped with substitutions that reduced the DUB activity but did not noticeably affect polyprotein cleavage (65). This is important because PL^pro^ plays a crucial role in virus replication during the liberation of nsp1 to nsp4 from pp1a and pp1ab. In earlier research, the impact of such DUB-inactivating substitutions was characterized in cell culture-based experiments in which mutant PL^pro^ domains were overexpressed and, in accordance with the results of others, were found to have lost the ability to suppress activation of the IFN-β/NF-κB promoter (65-67). Here we describe the generation of recombinant (r) DUB-negative MERS-CoVs with amino acid substitutions in PL^pro^, to analyse for the first time, the impact of the DUB function on infection. These mutant viruses had similar growth kinetics as wt rMERS-CoV, however mRNA levels for IFN-β and ISGs were significantly higher in cells infected with DUB-negative rMERS-CoV. In a lethal MERS-CoV infection model in mice, the DUB-negative virus was found to be strongly attenuated as the survival rate of mice infected with this mutant was significantly higher than for wt virus-infected mice. Furthermore, DUB-negative virus was cleared faster from the lungs and mRNA levels of IFNs, ISGs, and cytokines were upregulated earlier relative to those in control mice infected with wt rMERS-CoV. Collectively, this demonstrates that CoV PL^pro^’s DUB activity enhances virulence in an animal model and that selective removal of this activity could be a basis for the design of MLV vaccines.

## Materials and Methods

### Cell culture

HuH7 cells (a kind gift of Dr. Ralf Bartenschlager, Heidelberg University) were maintained at 37°C in 5% CO_2_ in Dulbecco’s modified Eagle’s medium (DMEM; Lonza) with 8% fetal calf serum (FCS; Bodinco BV), 100 units/ml penicillin (Lonza), 100 units/ml streptomycin (Lonza), 2 mM L-glutamine and non-essential amino acids (both PAA). MRC5 cells (ATCC CCL-171) were cultured in Eagle’s minimum essential medium (EMEM; Lonza) with the same supplements as used in the medium for HuH7 cells. BHK-21 cells (ATCC CCL-10) were grown in Glasgow minimum essential medium (Gibco) supplemented with 8% FCS, 100 units/ml penicillin, 100 units/ml streptomycin, 10% tryptose phosphate broth (Gibco), and 10 mM Hepes (pH 7.4; Lonza).

### Construction and launch of recombinant mutant MERS-CoV

A full-length MERS-CoV cDNA in a bacterial artificial chromosome (BAC) vector (68) was previously equipped with a T7 RNA polymerase promoter and a unique 3’-terminal NotI site for run-off transcription (43). After passaging of the MERS-CoV EMC/2012 isolate in Vero cells, a premature stop codon in ORF5 was observed in the minority of sequence reads, but this substitution became fixed upon additional virus passaging (4, 39). To limit the development of genetic variation in ORF5 upon passaging of rMERS-CoV in cells, we decided to introduce this premature stop codon at ORF5 codon 108 in the MERS-CoV full-length clone (further explanation on the use of this adaptation is provided in the Results section). This clone, pBAC-MERS-CoV-ORF5stop, was generated by two-step en-passant in vivo recombineering reactions in *E. coli* (69). Subsequently, substitutions in the PL^pro^-coding sequence were introduced in this clone using the same procedure. Primer sequences to introduce substitutions in pBAC-MERS-CoV are available upon request. Bacmids were isolated from bacteria and sequenced to verify the presence of the introduced substitution(s). pBAC-MERS-CoV-ORF5stop and derivatives thereof were linearized with *NotI* and they were purified by phenol-chloroform extraction and ethanol precipitation. Approximately 1 µg DNA was used as a template for *in vitro* transcription using the mMESSAGE mMACHINE T7 transcription kit (Thermo Fisher Scientific). BHK-21 cells (5×10^6^) were electroporated with 5 µg of this *in vitro* transcribed RNA using the Nucleofector 2b device (Lonza; program T-020) with Cell Line Nucleofector Kit T (Lonza). Immediately after electroporation, the cells were taken up in prewarmed cell culture medium and mixed with HuH7 cells (5×10^6^) before seeding in a 75-cm^2^ flask. Virus-containing supernatants were collected when complete cytopathogenic effect (CPE) was observed, normally 3 to 4 days after the electroporation. Harvested virus was passaged once in HuH7 cells to grow a virus stock used for further experiments.

### rMERS-CoV titration and sequencing

rMERS-CoV titers were determined by plaque assay on HuH7 or MRC5 cells, essentially as previously described for SARS-CoV (70). HuH7 cells were fixed after an incubation period of 3 days whereas MRC5 cells were fixed after 4 days. To determine the 50% tissue culture infectious dose (TCID50), 5-fold serial dilutions of pre-diluted rMERS-CoV stocks were prepared and used to infect HuH7 cells in flat-bottom microplates (96-wells). After 3 days, cells were fixed with formaldehyde, monitored for cytopathogenic effect, and the TCID50 was calculated using the Spearman & Kärber method (71).

In order to confirm the presence of the intended substitutions in the rMERS-CoV PL^pro^-coding sequence, RNA was isolated from virus-containing supernatants (passage 1) with the QIAamp Viral RNA Mini Kit (Qiagen). Total RNA was reverse-transcribed to cDNA using RevertAid H minus reverse transcriptase (Thermo Fisher Scientific) and random hexamers. The PL^pro^ domain (nucleotides 4435-5409 of MERS-CoV ORF1a/1ab) was amplified by PCR using Accuzyme DNA polymerase (Bioline) and after purification the PCR product was sequenced.

### Evaluation of rMERS-CoV replication and induced immune responses

HuH7 cells were infected with wt or mutant rMERS-CoV at a multiplicity of infection (MOI) of 0.01 or 5, to analyse multi- or single-cycle infections, respectively. MOI 0.01 or 1 was used to infect MRC5 cells. The rMERS-CoV inoculum was prepared in PBS containing DEAE (0.005% w/v) and 2% FCS, which was put on the cells for 1 h at 37°C. Inocula were removed and EMEM supplemented with antibiotics and 2% FCS was added to the cells. Following a high-MOI infection, cells were first washed three times with PBS before adding medium. Supernatants were harvested at various time points and rMERS-CoV titers were determined by plaque assay on HuH7 cells. After removal of the supernatant, infected cells were washed once with PBS and then lysed with RA1 buffer supplemented with β-mercaptoethanol for total RNA isolation using the NucleoSpin RNA II kit (Machery-Nagel). RNA served as a template in a reverse transcription (RT) reaction with RevertAid H minus reverse transcriptase and a combination of oligo(dT)_20_ primer (90%) and random hexamers (10%). To measure host immune responses, real-time quantitative (q) PCRs using iTaq SYBR Green Supermix (BioRad) were performed on a CFX384 Touch™ Real-Time PCR Detection System (BioRad). The following targets were amplified: IFN-β (5’-GTCACTGTGCCTGGACCATA-3’ and 5’-CTTGAAGCAATTGTCCCGT-3’), IFN-induced protein with tetratricopeptide repeats 2 (IFIT2; 5’-ATGTGCAACCTACTGGCCTAT-3’ and 5’-TGAGAGTCGGCCCATGTGATA-3’), viperin (5’-CGTGAGCATCGTGAGCAATG-3’ and 5’-TCTTCTTTCCTTGGCCACGG-3’), MERS-CoV genome (5’-CCCTCGACTCCAGGCTTCTGC-3’ and 5’-ATGTGCACACCGCGAGGCAT-3’), and ribosomal protein L13a (RPL13a; 5’-AAGGTGGTGGTCGTACGCTGTG-3’ and 5’-CGGGAAGGGTTGGTGTTCATCC-3’). Using

NormFinder software (72), RPL13a was identified as a suitable normalization gene. Quantification was done using the standard curve method using standard dilution series that were included in the qPCR. Furthermore, data was normalized against the relative quantities of RPL13a mRNA and mRNA levels of wt-infected MRC5 were set to 1. Significance relative to wt was calculated using an unpaired Student’s *t*-test and p values < 0.05 were considered statistically significant.

### rMERS-CoV infection of cytokeratin 18-*hDPP4* mice

Wild-type or V1691R rMERS-CoV grown on HuH7 cells (passage 1) was titrated by plaque assay on HuH7 cells as described above. Mice expressing hDPP4 under control of the cytokeratin (K) 18 promoter (73) were anesthetized with isofluorane and intranasally inoculated with MERS-CoV (1×10^5^ PFU per mouse) diluted in DMEM to a volume of 50 µl. Control mice were mock-infected with 50 µl DMEM. Mice were examined and weighed daily. The set endpoint was at 14 days post inoculation (p.i.), except for mice that died or reached a humane endpoint before that time. At other predetermined time points mice were euthanized, lungs were removed and divided for determination of the rMERS-CoV titers and for RNA isolation. In parallel, blood samples were taken and processed by centrifugation (1,000 x g for 5 min) to obtain sera, also before the rMERS-CoV infection sera was collected. Virus stocks used for the mouse experiment were prepared in the biosafety level 3 (BSL3) facilities at Leiden University Medical Center. The animal experiment itself was conducted in a BSL3 laboratory at Center for Animal Health Research (CISA-INIA) in Madrid, while sample analysis was performed at Leiden University Medical Center. Animal experimental protocols were approved by the Ethical Committee of the Center for Animal Health Research (CISA-INIA) (permit numbers: 2011–009 and 2011–09) in strict accordance with Spanish National Royal Decree (RD 1201/2005) and international EU guidelines 2010/63/UE about protection of animals used for experimentation and other scientific purposes and Spanish Animal Welfare Act 32/2007.

### rMERS-CoV isolation from K18-*hDPP4* mice

Lung parts from infected mice were weighed and placed in a gentleMACS M Tube (Miltenyi Biotec) containing 2 ml of PBS with 100 units/ml penicillin, 100 units/ml streptomycin (Lonza), 50 µg/ml gentamycin (Sigma-Aldrich), and 0.25 µg/ml Fungizone (Gibco). Lung tissues were homogenized with the gentleMACS dissociator by running program Lung_02 three times (Miltenyi Biotec). Homogenized samples were centrifuged at 9,500 x g for 5 min and supernatants were collected. Thereafter, rMERS-CoV titers were determined by plaque assay on HuH7 cells as described before. Statistical significance (p < 0.05) between wt and V1691R rMERS-CoV titers was determined using an unpaired Student’s *t*-test.

### RNA isolation from lungs of rMERS-CoV-infected mice and RT-qPCR

To preserve the integrity of RNA, lungs were soaked in RNAlater (Thermo Fischer Scientific) directly after harvesting, and stored at −80°C. For RNA extraction, lungs were transferred to a gentleMACS M Tube containing 2 ml of RA1 buffer (Machery-Nagel) supplemented with 1% β-mercaptoethanol. Tissue was dissociated in the gentleMACS dissociator using program RNA_02 (Miltenyi Biotec). Subsequently, samples were centrifuged at 9,500 x g for 5 min and supernatants were aliquoted. RNA was then isolated following the standard protocol of the NucleoSpin RNA II kit (Machery-Nagel). RT-qPCRs and their analysis were performed as described above but with the following deviation. To measure mRNA levels of CCL5, IL-6, and IFIT2, a triplex qPCR with TaqMan probes was performed with HotStarTaq Master mix (Qiagen) whereas all other targets were amplified using gene-specific primer sets and iTaq SYBR Green Supermix (BioRad). All primers and probe sequences can be requested.

The obtained cDNA was also used to verify that the V1691R substitution in MERS-CoV PL^pro^ was retained after several rounds of virus replication in mice. PCR and sequencing to determine the PL^pro^-coding sequence was executed as described above.

### Virus neutralization test

Serum dilutions were prepared in round-bottom microplates (96-wells) containing EMEM supplemented with antibiotics and 2% FCS. Serum samples were heat-inactivated at 56°C for 30 min, after which two-fold serial dilutions were prepared with a 1:5 dilution as starting point. To each well, an amount of MERS-CoV equalling 100 TCID50 units was added and incubated at 37°C for 1 h, to allow virus neutralization. Subsequently, samples (volume of 150 µl) were used to infect HuH7 cells (1×10^4^/0.32-cm^2^) which had been seeded in flat-bottom microplates (96-wells) one day earlier. Cells were incubated for 3 days at 37°C and subsequently fixed with the addition of 30 µl of 37% formaldehyde. Cell layers were microscopically assessed for CPE and subsequently stained with crystal violet. As a positive control for virus neutralization, rabbit antisera raised against a synthetic MERS-CoV antigen (S1-Fc, containing residues 358-588 of the viral S protein) were included (74). Back titration of MERS-CoV, to confirm the dose of the used inoculum was within the acceptable range of 30 to 300 TCID50, was done by adding two-fold dilutions starting with the diluted MERS-CoV (100 TCID50) to HuH7 cells. TCID50 was calculated using the Spearman & Kärber method (71). Virus neutralization tests were performed twice, and the mean of the two measurements was used.

## Results

### DUB-negative PL^pro^ mutants are viable and genetically stable

In our previous work, we described amino acid substitutions in the Ub-binding site of MERS-CoV PL^pro^ that specifically disrupt its DUB activity without affecting the overall proteolytic activity (65). Using ectopic expression of mutant PL^pro^ domains, we could implicate the DUB activity in antagonizing the host innate immune response (65). However, we now aimed to obtain direct proof that the enzyme also counteracts the type I IFN response in the context of the MERS-CoV-infected cell. A BAC-based MERS-CoV reverse genetics system (43, 68), based on the sequence of the EMC/2012 isolate (4), served as the starting point for this study.

A premature stop codon in ORF5 was seen in 45% of the sequence reads of the cell culture passage 6 MERS-CoV EMC/2012 isolate and appeared to be fixed after additional virus passaging in Vero cells (4, 39). Accessory ORFs like ORF5 are dispensable for MERS-CoV replication in cell culture and their protein products have been implicated in suppression of the innate immune response and NF-κB activation (45, 68). Indeed, deletion of ORF5 did not alter virus replication compared to wt MERS-CoV in Vero 81, HuH7, and Calu-3 cells (45, 75). However, higher levels of inflammatory cytokines were produced in Calu-3 cells upon infection with virus lacking ORF5 compared to wt virus (45). To avoid complications due to ORF5 evolution and associated changes in host immune suppression while generating PL^pro^ mutant virus stocks for our studies, all PL^pro^ mutants were engineered using a full-length construct containing the premature stop codon at ORF5 codon 108. Into this backbone, we subsequently introduced substitutions, based on the MERS-CoV pp1a/pp1ab amino acid numbering, V1691R, T1653R, and V1674S, as well as the combinations T1653R+V1691R and T1653R+V1674S+V1691R (named “double mutant” and “triple mutant”, respectively, from this point on). Val1691 and Thr1653 map to the S1 Ub-binding site of MERS-CoV PL^pro^ and have direct interactions with Ub, in contrast to Val1674 that was hypothesized to associate with the distal Ub of poly-Ub chains (65). In cell culture-based reporter assays, the V1691R substitution reduced the DUB activity of PL^pro^ most effectively, T1653R impaired it to a lesser extent, whereas V1674S only marginally reduced it (65). In all cases, proteolytic activity of the protease towards the polyprotein backbone remained intact.

To launch wt and mutant rMERS-CoV, BHK-21 cells were transfected with *in vitro* transcribed MERS-CoV RNA. Progeny virus was harvested and virus stocks were then grown in HuH7 cells. Virus-induced CPE was observed for the wt control as well as all DUB-negative mutants listed in the previous paragraph, indicating that these viruses were viable and cytopathogenic in cell culture. Virus titers were determined by plaque assay on HuH7 and MRC5 cells and additionally plaque phenotypes were examined. The plaque size of rMERS-CoV V1691R was slightly smaller compared to wt rMERS-CoV on both cell lines (**Fig 1**). Plaques of mutants T1653R and V1674S were comparable in size to those of the wt control. However, when combined with the V1691R substitution in double or triple mutants, plaque sizes were more variable and on average slightly decreased (**Fig 1**). To verify whether the substitutions were retained during virus propagation in HuH7 cells, RNA was isolated from virus-containing supernatant and used for RT-PCR amplification and sequencing. The PL^pro^-coding region of the genome was sequenced and after one passage the presence of all engineered substitutions was confirmed, in the absence of any additional (unintended) substitutions. Together, this demonstrated that rMERS-CoV mutants carrying changes in the Ub-binding site of PL^pro^ are viable and, at least initially, genetically stable in our experimental setting.

**Fig 1.**
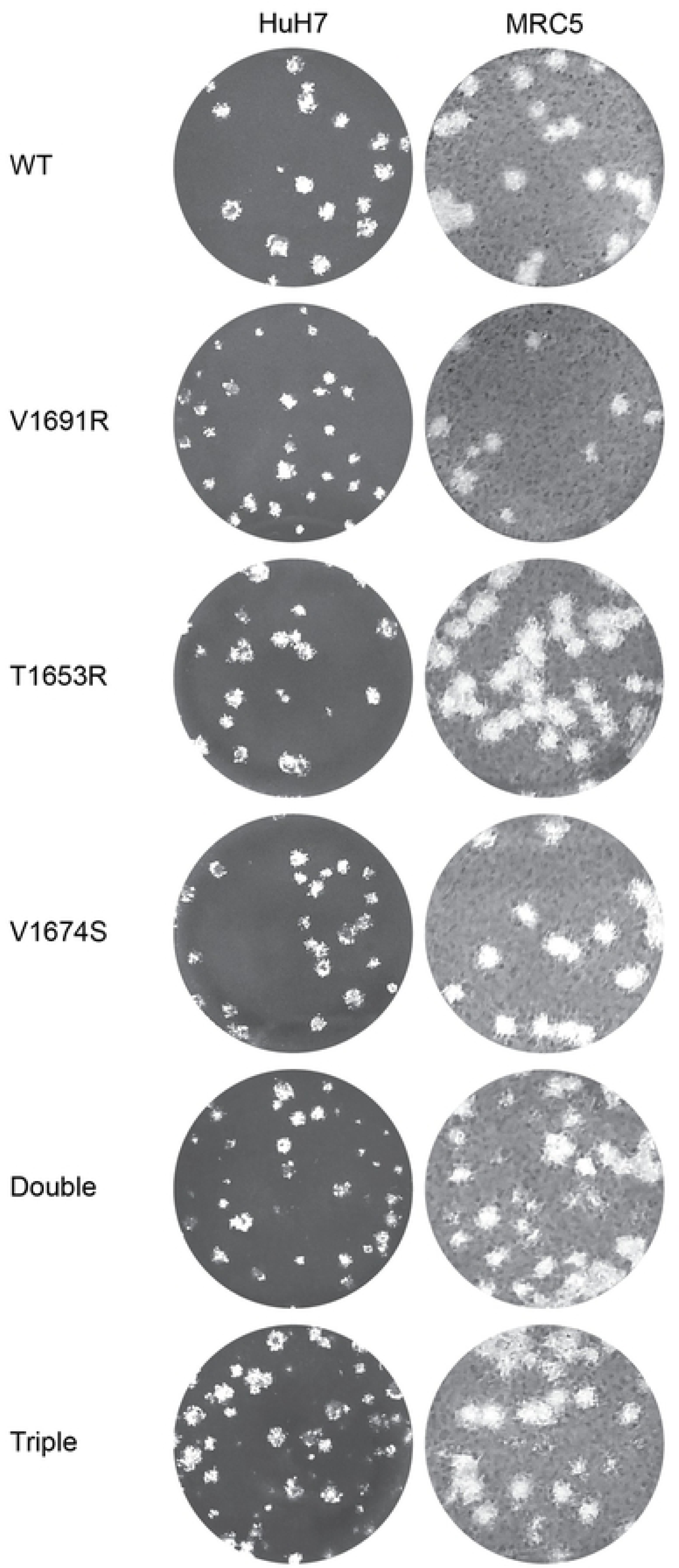
Plaque phenotype of rMERS-CoV PL^pro^ mutants. Plaque assays on HuH7 and MRC5 cells are presented for wt and DUB-negative rMERS-CoV with single, double or triple substitutions in PL^pro^.

### DUB-negative MERS-CoV PL^pro^ mutants have normal replication kinetics but induce an enhanced innate immune response

In order to characterize the rMERS-CoV PL^pro^ DUB-negative mutants, their replication kinetics and the induced innate immune responses were analyzed during single- and multi-cycle infections of HuH7 and MRC5 cells. For single-cycle experiments, we employed an MOI of 5 or 1 for HuH7 and MRC5 cells, respectively. V1691R, the single substitution that reduced PL^pro^’s DUB activity most effectively, when engineered into rMERS-CoV showed identical replication kinetics as wild-type rMERS-CoV, since the two viruses grew to similar titers in both cell lines at the different time points tested (**Fig 2A and B**). Also, the other mutants yielded titers comparable to wt virus 24 h p.i. when MRC5 cells were infected with MOI 1 (**Fig 2C**). Notably, rMERS-CoV peak titers were approximately 1-log higher in MRC5 cells compared to HuH7 cells. For multi-cycle infection experiments, cells were infected with an MOI of 0.01 and virus-containing supernatants were harvested 24 and 48 h p.i. At both these time points, there was no significant titer difference between wt virus and mutant V1691R (**Fig 2D-G**). Essentially similar results were obtained for all other DUB-negative mutants tested (**Fig 2D-G**), suggesting that overall replication in human cells is not affected by substitutions in the Ub-binding site of MERS-CoV PL^pro^.

**Fig 2.**
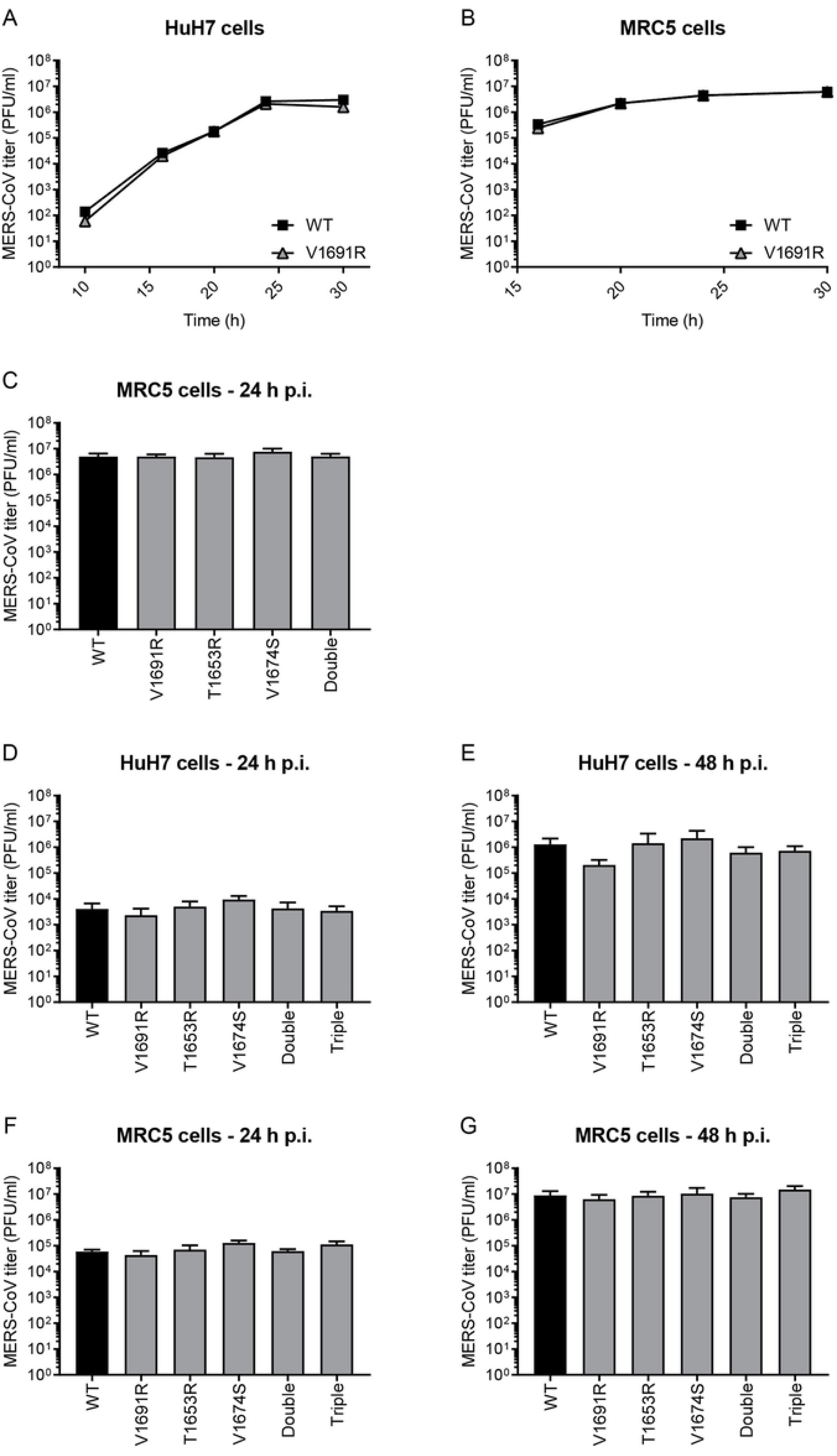
Growth of rMERS-CoV PL^pro^ mutants. HuH7 cells were infected with wt or DUB-negative rMERS-CoV with MOI 5 (A) or 0.01 (D and E), whereas an MOI of 1 (B and C) or 0.01 (F and G) was used to infect MRC5 cells. At different time points p.i., culture supernatants were harvested and rMERS-CoV progeny titers were determined by plaque assay on HuH7 cells. Growth curves (A and B) were performed once, while all other infections were repeated independently three times. Error bars represent the standard deviation.

Despite the lack of clear replication differences in cell culture, we anticipated that viruses with impaired DUB activity would be less capable of suppressing the innate immune response. Both MRC5 and HuH7 cells can mount an IFN response upon stimulation with poly(I:C), a synthetic mimic of dsRNA (data not shown). To assess whether DUB-negative mutants indeed induce a stronger innate immune response than wt rMERS-CoV, mRNA levels of IFN-β, IFIT2, and viperin in infected cells were measured by real-time RT-qPCR. Additionally, virus replication was evaluated by monitoring the intracellular accumulation of MERS-CoV genomic RNA. Data were normalized to house-keeping gene RPL13A, and are presented as fold increase over mock-infected cells. IFN-β levels were below the detection limit (Ct values > 35) in wt or DUB-negative rMERS-CoV-infected HuH7 cells, while these cells did have, although limited, upregulated IFN-β levels upon transfection of poly(I:C) (data not shown). Also in mock-infected MRC5 cells IFN-β mRNA was hardly detectable and therefore only a comparison was made between wt and DUB-negative-infected MRC5 cells. Upon single-cycle infection of MRC5 cells, significantly higher mRNA levels of IFN-β (∼4-fold), IFIT2 (∼3-fold), and viperin (∼4-fold) were measured upon infection with V1691R rMERS-CoV compared to wt rMERS-CoV (**Fig 3A-C**). The double mutant showed a more pronounced increase while the single T1653R mutation did not affect the mRNA levels of the qPCR targets at all (**Fig 3A-C**). Also, cells infected with viruses carrying the V1674S substitution induced a very low response comparable to that against wt virus (**Fig 3A-C**). Wild-type and mutant viral genomic RNA levels were similar in infected cells (**Fig 3D**). In multicycle infections of MRC5 cells, V1691R rMERS-CoV induced higher levels of IFN-β and ISGs mRNA than wt virus (IFN-β ∼3-fold, IFIT2 ∼4-fold, and viperin ∼8-fold; **Fig 3E-G**). This was even more pronounced with the double mutant, while addition of the V1674S substitution unexpectedly reduced IFN-β and ISG responses relative to the double mutant (**Fig 3E-G**). The additive effect of T1653R implies that this substitution, when combined with V1691R, is further reducing Ub-binding. PL^pro^ mutants T1653R and V1674S generated responses comparable to those against wt virus, and virus genome levels were similar (**Fig 3E-H**). Interestingly, MERS-CoV genome levels were lower in cells infected with V1691R, with the double, and with the triple mutant, compared to wt virus in these multicycle infections (**Fig 3H**), indicating that these mutations eventually do have impact of the replication of the virus, probably due to the higher innate immune responses they elicit. Collectively, viruses containing the T1653R or V1674S substitution did not induce different host responses in MRC5 cells compared to wt virus. In contrast, V1691R rMERS-CoV elicited a clear upregulation of IFN-β and ISG mRNAs, which was even more prominent when combined with the T1653R substitution.

**Fig 3.**
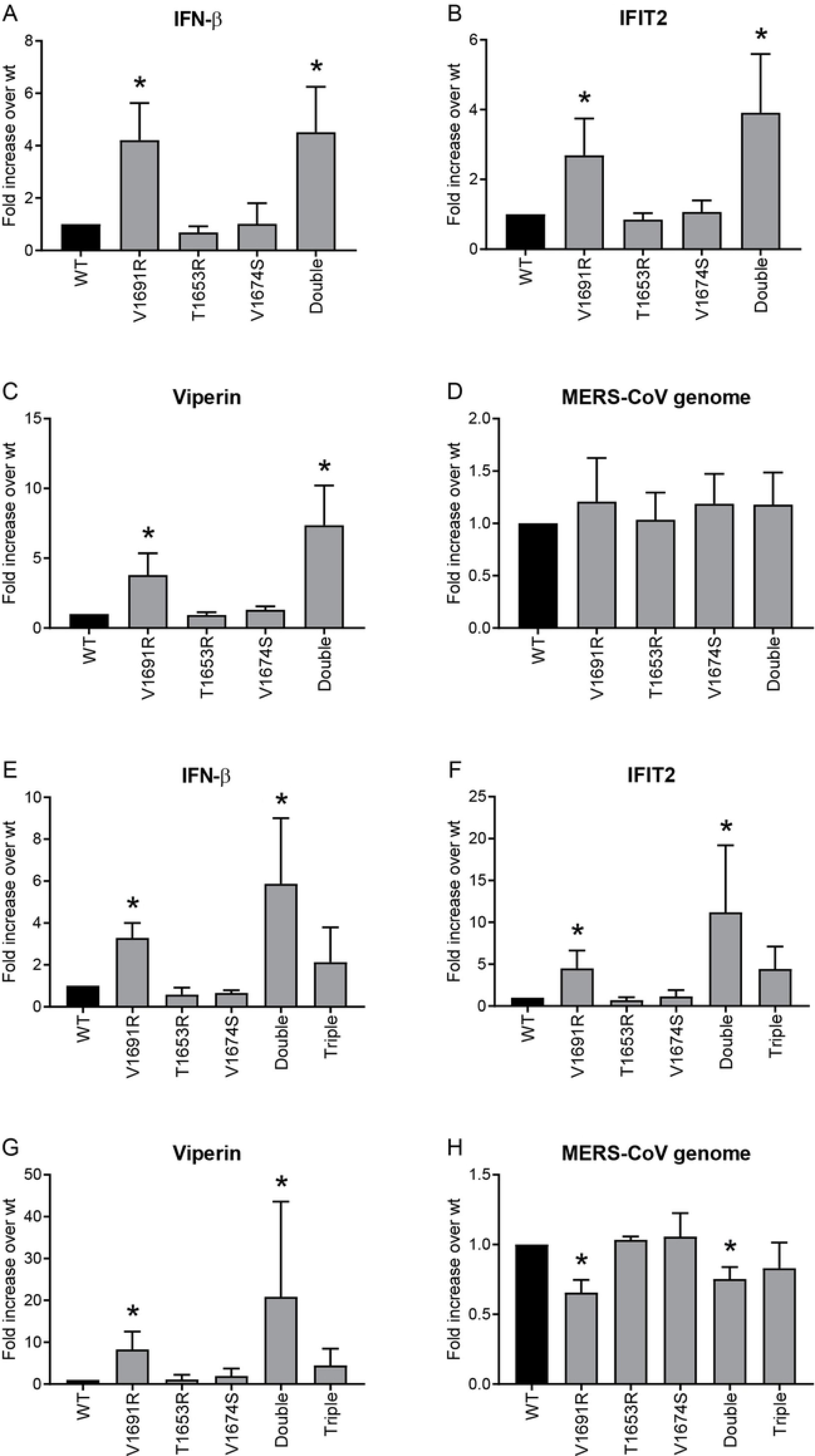
Immune responses in MRC5 cells infected with wt or DUB-negative rMERS-CoV. MRC5 cells were infected with wt or DUB-negative rMERS-CoV with MOI 1 (A-D) or 0.01 (E-H) and intracellular RNA was isolated at 24 h or 48 h p.i., respectively. Levels of IFN-β mRNA (A and E), IFIT2 mRNA (B and F), viperin mRNA (C and G), and MERS-CoV genomic RNA (D and H) were determined by real-time RT-qPCRs and normalized to transcript levels of RPL13a. Results were obtained in three independent experiments, error bars represent the standard deviation, and asterisks show a significant difference in elicited responses between wt and DUB-negative rMERS-CoV (p < 0.05).

### Increased survival of mice infected with DUB-negative rMERS-CoV

As shown in **Fig 2**, rMERS-CoV V1691R and wt virus had similar replication kinetics in mammalian cells, but infection with the former resulted in an upregulation of the mRNA levels for IFN-β and ISGs (**Fig 3**). This prompted us to evaluate this DUB-negative virus in a lethal mouse model for MERS-CoV infection, which relies on transgenic mice expressing the human DPP4 (hDPP4) receptor under the control of the K18 promoter (73). Intranasal inoculation of these mice leads to MERS-CoV replication in the lungs and production of interferons, cytokines, and chemokines. Ultimately, the animals develop pathological changes in their lungs but also in their brain, and most likely the mice die of neurological disease due to MERS-CoV infection in the brain (73).

We aimed to investigate V1691R rMERS-CoV infection of these transgenic mice with respect to its lethality and the innate immune responses in the lung. To this end, mice were infected intranasally with 1×10^5^ PFU of either wt or DUB-negative rMERS-CoV, whereas a third group of mice was mock-infected. Each group contained a total of 16 mice of which 12 animals per group were euthanized at early time points. The remaining four animals from each of these three groups were monitored for a period of 14 days to assess their survival and measure their body weight on a daily basis. All four mice infected with wt rMERS-CoV died or had to be euthanized after reaching a humane endpoint by day 7 or 8 p.i. (**Fig 4A**). In contrast, only one of the four mice infected with DUB-negative rMERS-CoV died at day 8 p.i., while the other three mice survived for the full 14-day period (**Fig 4A**). Even with the small number of animals used for this experiment, this difference in survival between mice infected with wt or DUB-negative rMERS-CoV is statistically significant (log-rank test p value = 0.028). Both wt and DUB-negative rMERS-CoV-infected mice lost weight over the course of infection (**Fig 4B**), but in all three surviving mice infected with DUB-negative rMERS-CoV this weight loss was reversed from day 10 p.i. forward (**Fig 4C**). The results clearly suggested that the virulence of the DUB-negative virus was reduced compared to wt virus.

**Fig 4.**
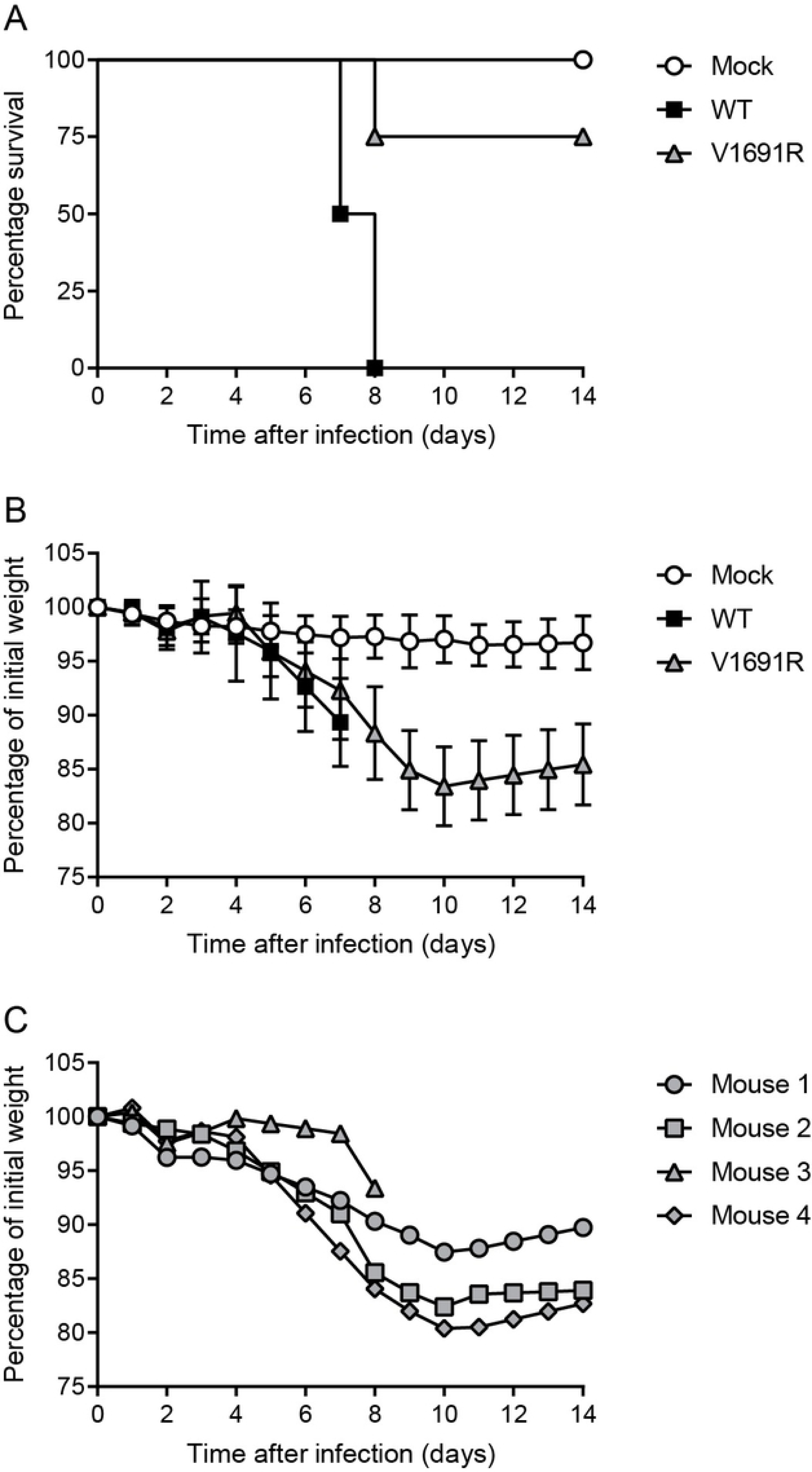
Survival and weight loss of mice infected with wt or DUB-negative rMERS-CoV. Mice that express hDPP4 under the control of the K18 promoter were infected with 1×10^5^ PFU wt or DUB-negative V1691R rMERS-CoV, or were mock-infected. Survival data (A) are from four infected animals per group. The weight of the mice (B) was monitored daily and the mean of the group of four mice is presented with error bars showing the standard deviation. The weight of each individual mouse infected with DUB-negative rMERS-CoV is presented in (C).

### DUB-negative rMERS-CoV is cleared faster from mice lungs than wt virus

To measure rMERS-CoV replication and the innate immune responses the viruses elicited in lungs of infected mice, we euthanized four animals from each group at days 1, 2, and 4 p.i. Lungs were removed and part of the material was homogenized and used for virus titration. For wt rMERS-CoV, progeny titers were approximately 1×10^6^ PFU per g of lung tissue during the first four days after infection (**Fig 5**). The DUB-negative virus reached similar virus titers as the wt control during the first two days p.i., indicating that this virus also replicates well in mice. Unexpectedly, however, it proved impossible to recover virus from the lungs of three of the mice infected with DUB-negative rMERS-CoV (twice in the day 1 p.i. group, and once in the day 2 p.i. group; **Fig 5**). Consequently, it remains unclear whether these three animals had been successfully inoculated. All DUB-negative rMERS-CoV-infected mice that were euthanized at day 4 p.i. were virus-positive, and their lung titers were significantly lower than in wt virus-infected mice at that time point (**Fig 5**), suggesting that the DUB-negative virus is cleared faster from the lungs than wt rMERS-CoV. Virus titers were also determined in mice that died from the infection at day 7-8 p.i. and in survivors of the infection at day 14. By day 7-8 p.i., the rMERS-CoV wt titers were around 1×10^4^ PFU per g of lung tissue, signifying that virus titers decreased over time (**Fig 5**). The virus titer of the single mouse that died of the DUB-negative virus infection was 2×10^3^ PFU per g of lung tissue (**Fig 5**). Virus could not be recovered from the lungs of any of the mice that had survived the infection with DUB-negative rMERS-CoV after 14 days (**Fig 5**).

**Fig 5.**
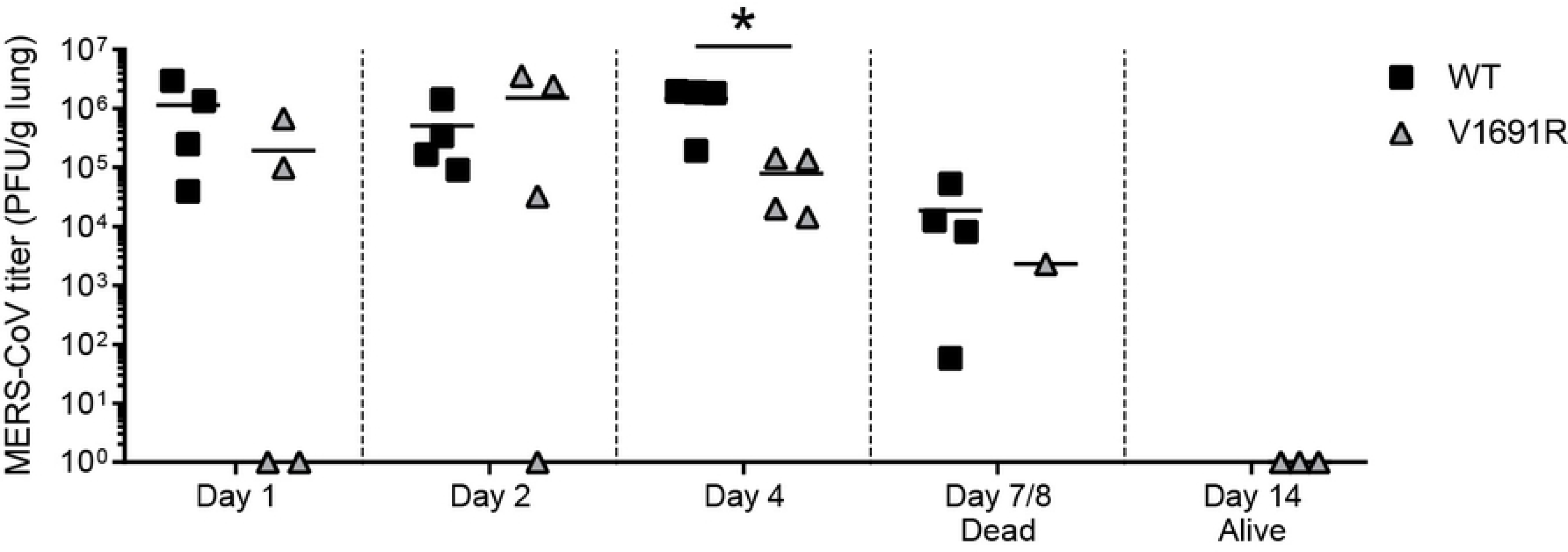
Replication of wt and DUB-negative rMERS-CoV in mice. Groups of four K18-*hDPP4* mice infected with wt or V1691R rMERS-CoV were euthanized at days 1, 2, or 4 after infection. Lungs were removed and homogenized, and rMERS-CoV lung titers were measured by plaque assay on HuH7 cells. Recombinant MERS-CoV titers in lungs were also determined in mice that died at day 7 or 8 after the infection, and in the three DUB-negative rMERS-CoV-infected mice that were euthanized at 14 days p.i. The mean per group and the rMERS-CoV titer in PFU per gram of lung tissue are presented for each mouse. The asterisk indicates a significant difference between wt and V1691R rMERS-CoV, determined with a Student’s *t*-test (p < 0.05).

In order to verify whether the DUB-inactivating substitution in the MERS-CoV mutant was stably maintained during the animal experiment, sequencing of the PL^pro^-coding region of the genome was performed. To this end, lung tissue was homogenized, RNA was isolated, and the PL^pro^-coding sequence was amplified by RT-PCR and sequenced. In wt virus-infected mice, the consensus sequence of the PCR product was found to be identical to the sequence of the BAC-based cDNA clone from which the rMERS-CoV had been launched (data not shown). RT-PCR and sequencing were successful using RNA isolated from all mice infected with DUB-negative rMERS-CoV from which also DUB-negative rMERS-CoV was recovered (**Fig 5**), and revealed the presence of the V1691R substitution in PL^pro^ without additional substitutions in PL^pro^ (data not shown). No virus-specific RT-PCR product could be obtained from the lungs of mice that were found negative for infectious virus at day 1 or 14 p.i. (data not shown; **Fig 5**). For one mouse inoculated with DUB-negative rMERS-CoV, although virus could not be isolated at 2 days p.i. (**Fig 5**), a virus-specific RT-PCR product could be obtained using another piece of its lung, after which the presence of the PL^pro^ mutation could be confirmed by sequencing. Possibly, in this particular mouse, the virus was distributed unequally in the lungs during intranasal inoculation, explaining why only some parts were virus-positive at 2 days p.i. Similar issues may explain the lack of virus isolation for the two mice that were negative for the mutant virus at 1 day p.i., as for the described experiments not the entire lungs were used, and as there was no obvious technical reason to suggest inoculation failure. Overall, our data established that the DUB-negative virus, which stably maintained the V1691R substitution, is cleared faster from the lungs of infected mice than wt virus.

### Accelerated innate immune responses following infection with DUB-negative rMERS-CoV

We hypothesized that the significantly higher survival rate of mice infected with the V1691R mutant and the faster clearance of DUB-negative rMERS-CoV from the lungs were the result of enhanced innate immune responses due to the loss of PL^pro^’s DUB activity. To investigate this, we analyzed the mRNA levels of innate immune factors and cytokines from the lungs of infected mice. Real-time RT-qPCRs were performed and results were normalized to the mRNA of RPL13A, a housekeeping gene selected for its stability in the infected mice. The selected mRNA targets were representative for innate immune responses that are relevant for MERS-CoV infection and included IFNs (IFN-β and IFN-λ), ISGs (ISG15 and IFIT2), the cytokine TNFα that is controlled by NF-κB activation, the pro-inflammatory cytokine IL-6, and the chemokine CCL5 (**Fig 6**) (76).

**Fig 6.**
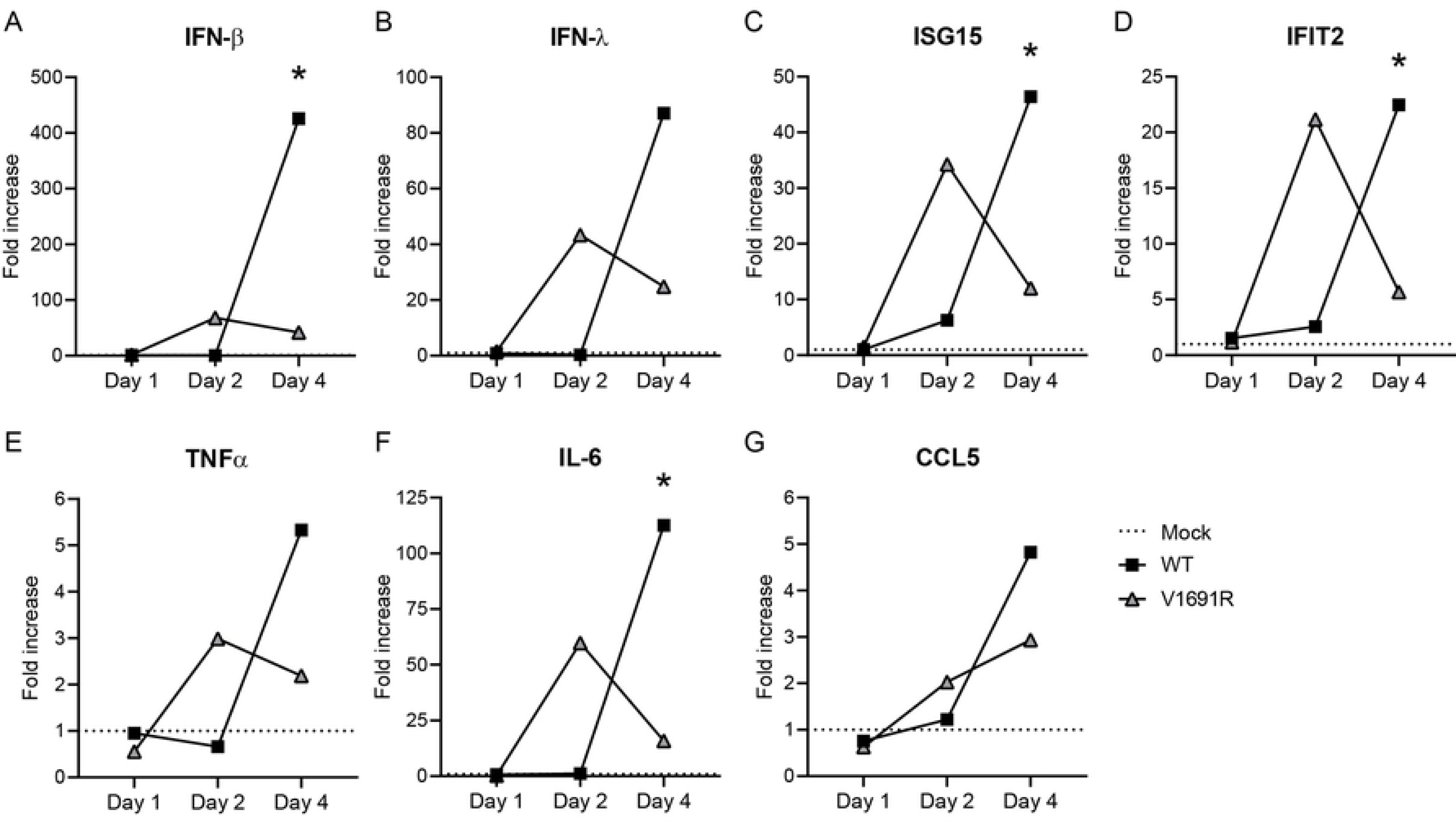
Innate immune responses of mice infected with wt or DUB-negative rMERS-CoV. RNA was isolated from lungs of mice infected with wt or DUB-negative rMERS-CoV, or mice that had been mock-infected. RNA was used in real-time RT-qPCRs to measure the fold induction of the mRNA levels of IFN-β (A), IFN-λ (B), ISG15 (C), IFIT2 (D), TNFα (E), IL-6 (F) and CCL5 (G). Results were analysed using the standard curve method and normalised to the mRNA level of the RPL13a gene. The mean per group is presented and the dashed line represents the average that was set to 1 for the mock-infected mice. Statistical significance between wt and DUB-negative MERS-CoV-infected mice is shown (* p < 0.05).

After the first day of infection, the mRNA levels for all targets were similarly low in wt and DUB-negative rMERS-CoV-infected mice (**Fig 6**). At 2 days p.i., transcript levels in wt rMERS-CoV-infected mice were comparable to those in mock-infected mice while a clear increase was measured in DUB-negative rMERS-CoV-infected mice (**Fig 6**). For ease of visualization, the mean fold increase for each target is presented in **Fig 6**, while the fold increase for each individual mouse is displayed in **S1 Fig** to be able to show the variation in the responses between mice in the same group. Notably, the IFN-β and IFN-λ values for the wt MERS-CoV-infected mice showed less variation than those of the mock-infected animals, suggesting that the wt virus efficiently suppressed the expression of these cytokines at day 2 p.i. (**S1A and B Fig,** compare fold increase of mock and wt-infected mice at day 2 p.i.). The mean level of IFN transcripts in DUB-negative rMERS-CoV-infected mice was clearly, although not statistically significantly, higher than in the mock and wt virus-infected mice (**Fig 6A and B**). ISG15 and IFIT2 mRNA levels were found to be slightly elevated in wt rMERS-CoV-infected mice relative to mock-infected mice, while an even more pronounced increase was seen in DUB-negative rMERS-CoV-infected mice (**Fig 6C and D**). The upregulation of IFN-β mRNA in individual DUB-negative MERS-CoV-infected mice corresponded to the increased levels of ISG mRNAs in the same mice at 2 days p.i. (**S2 Fig**) This illustrates that the ISG responses measured were likely induced by upregulation of type I IFNs in these animals. Due to the limited group size and considerable variation in the responses between the four mice that received DUB-negative rMERS-CoV (**S1 Fig**, see fold increase of DUB-negative virus-infected mice at day 2 p.i.), the difference in mRNA levels between wt and DUB-negative rMERS-CoV-infected mice at 2 days p.i. was not statistically significant. However, the trend of an accelerated increase of mRNA levels was clearly there for all innate immune targets upon infection with DUB-negative rMERS-CoV. This strongly suggests that, compared to the wt control virus, the DUB-negative virus is less able to suppress innate immune responses resulting in an earlier reaction.

By day 4 p.i., a very different innate immune response in the mice was seen compared to the response at 2 days p.i. (**Fig 6**). In the DUB-negative virus group, the increase of innate immune transcripts was slightly lower relative to 2 days p.i. (**Fig 6**), and the variation in mRNA levels between the four mice was smaller in this group at 4 days than at 2 days p.i. (**S1 Fig**, compare e.g. the IFN-β fold increase of each individual V1691R-infected mouse at day 2 p.i. with day 4 p.i.), suggesting down-regulation of the response by day 4 p.i. in this group of mice. Wild-type virus-infected mice showed, however, a very strong increase in transcript levels for all tested targets between 2 and 4 days p.i. Especially the mRNA levels of IFN-β, IFN-λ, and IL-6 were remarkably upregulated (**Fig 6A, B, and F**). Together, this demonstrates that between 2 and 4 days p.i. the mRNA levels of innate immune factors, cytokines, and chemokines were already down-regulated in the DUB-negative virus-infected mice, in contrast to the strong upregulation observed for rMERS-CoV wt-infected mice.

We expected that the early immune responses would be further down-regulated over time in the mice infected with the DUB-negative mutant. To confirm this, mRNA levels were determined in lungs of the three surviving mice infected with V1691R rMERS-CoV and mock-infected mice at 14 days p.i. The number of transcripts of innate immune factors and cytokines was similar between these two groups, indicating that all mRNAs in surviving animals were back to an almost basal level (**S3 Fig**). In summary, DUB-negative MERS-CoV induced an earlier, and subsequently again downregulated, innate immune response, whereas wt virus-infected mice show a relatively late and excessive response.

### Neutralizing antibodies in DUB-negative rMERS-CoV-infected mice

The three mice that survived the infection with DUB-negative mutant rMERS-CoV V169R were euthanized 14 days p.i. and their sera were collected. Prior to infection, their pre-immune sera had also been obtained. To determine whether a neutralizing antibody response against MERS-CoV had developed, despite the relatively recent infection, we performed virus neutralization tests (VNTs). Pre-immune sera did not contain antibodies that could neutralize MERS-CoV (VNT titer <5) (data not shown). This was also true for the day 14 p.i. serum of one mouse. However, in the two other mice low levels of virus-neutralizing antibodies were measured (VNT titers of 15 and 20), suggesting that their adaptive immune system had been activated by the infection with DUB-negative rMERS-CoV, resulting in the production of antibodies that have neutralizing capacity.

## Discussion

Many viral proteases with a primary role in virus replication have evolved to target host cell proteins, and often these accessory cleavage events indirectly affect viral replication and pathogenesis (77). Coronaviral PL^pro^s constitute a striking example of this kind of dual functionality, as they not only cleave multiple sites in the replicase polyproteins of which they are part, but can also deconjugate Ub, and Ub-like modifiers such as ISG15, presumably to suppress innate immune responses (55). Studying this DUB activity in the context of the CoV-infected cell is difficult because both PL^pro^ functionalities rely on the same enzymatic active site, which cannot be inactivated without killing virus replication. Therefore, in our earlier research, we explored an alternative approach to selectively inactivate the DUB activity of MERS-CoV PL^pro^. Based on the available crystal structure of the PL^pro^-Ub complex, we disrupted the enzyme’s Ub-binding site to minimize its affinity for Ub (65). Based on assays carried out in expression systems, it was concluded that the DUB activity of PL^pro^ could indeed be inactivated almost completely without affecting the proteolytic activity needed for viral polyprotein cleavage (65). Subsequently, our study suggested that the DUB activity of MERS-CoV PL^pro^ may indeed be involved in suppressing innate immune responses, as DUB-negative PL^pro^ was no longer able to downregulate IFN-β promoter activity. We have now engineered mutant DUB- negative viruses, thereby convincingly extending these observations to MERS-CoV infected-cells, in culture and *in vivo*. We characterized the replication and the innate immune responses these viruses induce in comparison to wt virus.

Interestingly, some of the DUB-negative viruses, specifically those containing the V1691R substitution, induced higher levels of IFN-β and ISG mRNAs than wt virus in MRC5 cells, confirming a role of PL^pro^ in innate immune suppression, now demonstrated during infection (**Fig 3**). However, this upregulation of the type I IFN response did not reduce the progeny titers of DUB-negative rMERS-CoV in MRC5 cells (**Fig 2**). We did observe that the plaque size of rMERS-CoV containing the V1691R substitution was slightly smaller when compared to wt virus (**Fig 1**), suggesting that the mutant virus was somehow affected after all. Also, in multi-cycle infections, MERS-CoV genome levels were reduced upon infection with DUB-negative MERS-CoVs carrying the V1691R substitution (**Fig 3H**), confirming a modest effect on virus RNA replication in cell culture. This suggests that, in this experimental set-up, the increased IFN expression and subsequent ISG response somewhat influences viral RNA replication but not, at least not noticeably, the production of infectious progeny. The decrease in MERS-CoV genome RNA levels was not observed in single-cycle infections in cell culture, highlighting that primary replication of the mutant viruses is not affected by the substitution in PL^pro^. In summary, by specifically disrupting the PL^pro^ interaction surface that binds Ub, we have demonstrated that the DUB activity of PL^pro^ indeed contributes to innate immune evasion in the context of the MERS-CoV-infected cell.

The importance of vDUB activity in the context of infection has thus far been tested for only a few viruses. Previously, the vDUB activity of equine arteritis virus (EAV), belonging to the order *Nidovirales* like CoVs, and Crimean-Congo haemorrhagic fever virus (CCHFV), a nairovirus in the order *Bunyavirales*, were specifically eliminated by mutagenesis (78, 79). These vDUBs are members of the ovarian tumor domain-containing family of DUBs and are structurally different from CoV PL^pro^s, which belong to the ubiquitin-specific protease DUB family. The replication of CCHFV was attenuated in IFN-competent cells upon DUB inactivation, due to the induction of a more robust immune response (78). DUB-negative EAV replicated like wt virus in equine lung fibroblasts, but did elicit enhanced innate immune responses (79). Nevertheless, *in vivo* the mutant virus provided an equal level of protection as the parental (attenuated) virus (80). Recently, the structural equivalent of the MERS-CoV V1691R substitution was introduced into SARS-CoV PL^pro^, yielding mutant SARS-CoV with a M1748R substitution (using SARS-CoV pp1a/pp1ab amino acid numbering) (81). Compared to the wt control, replication of this mutant virus was slightly delayed in cell lines that can mount an innate immune response, and it induced higher IFN-β mRNA levels (81). We here took the characterization of DUB-negative CoV mutants to the next level, by testing a DUB-negative MERS-CoV mutant in an animal model for the first time.

Recombinant MERS-CoV V1691R was tested in a transgenic mouse model for lethal MERS-CoV infection, which is based on hDPP4 expression under control of the K18 promoter (73). Consequently, hDPP4 was expressed in the lungs, brain, and various other organs. After intranasal inoculation, MERS-CoV replication was observed in lungs and wt rMERS-CoV titers were stable over the first four days of infection, showing 1×10^6^ PFU per g of lung tissue (**Fig 5**). Although this is lower than the previously reported peak titer at 2 days p.i. of 6×10^7^ PFU per g of lung tissue (73), it shows that we obtained robust virus replication of rMERS-CoV in these mice. In the previous study by Li *et al*., mice died at 6-7 days p.i. most likely due to neurological disease as a result of the infection in the brain in which MERS-CoV titers increased over time, although a contribution of lung damage to the eventual cause of death may also be likely (73). In wt MERS-CoV-infected K18-*hDDP4* mice lung pathology was indeed observed, characterized by vascular thrombi, alveolar oedema, degenerating cells, and cellular debris in lymphatic vessels (73). In accordance with those data, wt rMERS-CoV infection was lethal also in our hands, and all mice died at 7-8 days after infection (**Fig 4A**). Likely, the mice that succumbed to the infection also had MERS-CoV infection of the brain, however, the experimental circumstances did not allow us to take samples to support this assumption. Strikingly, infection with the DUB-negative virus resulted in a significantly increased survival compared to the animals infected with the wt virus (**Fig 4A**). The DUB-negative virus reached the same titers in the lungs as the wt virus by 2 days p.i., but these had dropped significantly by 4 days p.i. while they stayed high in the wt virus-infected mice (**Fig 5**). In future research, it will be interesting to determine whether histopathology is reduced in mice infected with DUB-negative rMERS-CoV to confirm that earlier virus clearance and reduced tissue damage correlates with increased survival, relative to in wt virus-infected mice.

Messenger RNA levels for IFNs (IFN-β and IFN-λ), ISGs (ISG15 and IFIT2), and pro-inflammatory cytokines (IL-6 and TNFα) started to rise in lungs of DUB-negative rMERS-CoV-infected mice at 2 days p.i. (**Fig 6**). Although there was no statistically significant difference compared to wt virus-infected mice, the trend of faster upregulation of these mRNAs was quite clear. Moreover, the fact that the levels for all these transcripts increased simultaneously strengthens the conclusion that the DUB-negative rMERS-CoV induced an accelerated innate immune response compared to the wt virus, again confirming the role of the PL^pro^ DUB activity in suppression of innate immune responses, now also in the context of a mouse model. The transcript levels of IFNs and cytokines slightly decreased in DUB-negative rMERS-CoV-infected mice at 4 days p.i., but in wt rMERS-CoV-infected mice the mRNA levels were extremely upregulated (**Fig 6**). We therefore hypothesize that the difference in survival rate is likely due to an accelerated, and after induction properly down-regulated, innate immune response in DUB-negative virus-infected mice, which is likely more effectively clearing the virus and preventing tissue damage, protecting these mice from lethality (**Figs 4-6**). In contrast, the response in wt virus-infected mice is first suppressed and then at later time points it is extremely upregulated (**Fig 6**). These results resemble the late and excessive IFN and pro-inflammatory cytokine response that likely contributes to lung immunopathology as observed for SARS-CoV infection in mice (38, 76). Recently, it was also demonstrated that the timing of the type I IFN response was crucial in the outcome of a MERS-CoV infection in mice (82). In line with our results, early administration of type I IFN was protective (comparable to responses induced by DUB-negative rMERS-CoV) whereas late administration promoted lethality (like in wt rMERS-CoV-infected mice) (82). Together, these observations support a concept in which delayed and aberrantly regulated innate immune response as a result of viral innate immune evasion importantly contributes to pathogenesis during infection. Notably, it is possible that other cellular processes besides innate immunity are influenced by PL^pro^s DUB activity, which might also affect mice survival. Therefore, it remains important to make a full inventory of cellular protein targets that are deubiquitinated by PL^pro^, and to assess how such targets affect host responses and survival by influencing the innate immune system and other cellular processes. Thus, the DUB-negative rMERS-CoV can be an important tool to further explore CoV-host interactions.

In two of the mice that survived the DUB-negative rMERS-CoV infection, we could measure neutralizing antibodies against MERS-CoV. The third surviving mouse did, at this relatively early time point of 14 days p.i., not have a detectable neutralizing antibody titer. The observed variation might be related to the course of infection, as the weight loss of this third mouse was the smallest of the three mice that survived the infection (**Fig 4C**, mouse 1), suggesting that MERS-CoV infection was milder than in the other two animals. It will be interesting to measure the acquired protection in the DUB-negative MERS-CoV infected mice by challenging them with (lethal) MERS-CoV. Although more in-depth characterization is necessary, the DUB-negative MERS-CoV may be a MLV vaccine candidate that can induce antibody responses as well as T-cell responses, which both are important for providing long-term effective protection against CoV infections (83-85). Interestingly, for MERS-CoV, vaccines are currently being designed for use in humans as well as dromedary camels, the latter aiming to reduce virus circulation and transmission to humans (84-86).

In summary, our results demonstrate, for the first time, that the presence of PL^pro^’s DUB activity causes a delay in the innate immune response, which has a major impact on pathogenesis in the mouse model used. Pathogenesis and lethality can thus be reduced by removal of the DUB activity, resulting in earlier, better-regulated and therefore likely more effective innate immune response that helps to clear the virus and decreases pathogenicity, despite the initial viral replication being similar to that seen in wt virus infections. Our study has thus also provided the proof-of-concept for the *in vivo* attenuation of MERS-CoV, which provides a basis to advance the analysis of the innate response as well as antibody and T-cell-mediated responses triggered by infection with DUB-negative MERS-CoV. It creates the opportunity to apply this strategy to develop MLV vaccines against MERS-CoV as well as other CoVs of societal or economic importance, such as porcine epidemic diarrhoea virus, feline CoV, avian infectious bronchitis virus, and bovine CoV. Also in the CoV field, the “attenuation-by-design” approach to eliminate one or more innate immune evasive functions may offer a general and effective strategy for MLV development.

## Acknowledgements

We are grateful to Dr. Jelle Goeman (Leiden University Medical Center) for assisting with the statistical analysis of the mouse experiment. We kindly thank Dr. Clara C. Posthuma and Jessika C. Zevenhoven-Dobbe (both Leiden University Medical Center) for initial modifications to pBAC-MERS-CoV. We are grateful to Irina C. Albulescu (Leiden University Medical Center) for developing the VNT for MERS-CoV and to Dr. Els van der Meijden (Leiden University Medical Center) for setting up the triplex RT-qPCR for monitoring mouse innate immune response. We thank Dr. Ralf Bartenschlager (Heidelberg University) and Dr. Berend Jan Bosch (Utrecht University) for providing reagents. We are grateful for regular inspiring discussions with Dr. Ben A. Bailey-Elkin and Dr. Brian L. Mark (both University of Manitoba).

## Supporting information

**S1 Fig. Innate immune responses of each infected mouse.** In the lungs of wt virus, DUB-negative virus, or mock-infected mice, the fold induction in mRNA levels of IFN-β (A), IFN-λ (B), ISG15 (C), IFIT2 (D), TNFα (E), IL-6 (F) and CCL5 (G) was measured by real-time RT-qPCR. The mean per group as well as the results per individual mouse are presented, and statistical significance is shown (* p < 0.05).

**S2 Fig. Innate immune responses of individual rMERS-CoV-infected mice.** Real-time RT-qPCR were performed on RNA isolated from wt or V1691R rMERS-CoV-infected mice at 2 days p.i. mRNA levels of IFN-β, ISG15 and IFIT2 for each individual mouse are shown, the dashed line at a fold increase of 1 indicates the set average of mock-infected mice.

**S3 Fig. Immune responses in mice that survived the infection with DUB-negative rMERS-CoV.** Isolated RNA from lungs of mice infected with DUB-negative rMERS-CoV or mice that were mock-infected was used in real-time RT-qPCRs. mRNA levels of IFN-β (A), IFN-λ (B), ISG15 (C), IFIT2 (D), TNFα (E), IL-6 (F) and CCL5 (G) were analyzed using the standard curve method and were normalized to relative levels of RPL13a. The mean and the value of each individual mouse is presented.

